# Elucidating the Membrane Binding Process of a Disordered Protein: Dynamic Interplay of Anionic Lipids and the Polybasic Region

**DOI:** 10.1101/2023.08.02.551595

**Authors:** Azadeh Alavizargar, Maximilian Gass, Michael P. Krahn, Andreas Heuer

## Abstract

Intrinsically disordered regions of proteins are responsible for many biological processes such as in the case of liver kinase LKB1 – a serine/threonine kinase, relevant for cell proliferation and cell polarity. LKB1 itself becomes fully activated upon recruitment to the plasma membrane by binding of its disordered C-terminal polybasic motif consisting of eight lysines/arginines to phospholipids. Here we present extensive molecular dynamics (MD) simulations of the polybasic motif interacting with a model membrane composed of phosphatidylcholin (POPC) and phosphatidic acid (PA) and cell culture experiments. Protein-membrane binding effects are due to the electrostatic interactions between the polybasic amino acids and PAs. For significant binding the first three lysines turn out to be dispensable, which was also recapitulated in cell culture using transfected GFP-LKB1 variants. LKB1-membrane binding results in a non-monotonous changes in the structure of the protein as well as of the membrane, in particular accumulation of PAs and reduced thickness at the protein-membrane contact area. The protein-lipid binding turns out to be highly dynamic due to an interplay of PA-PA repulsion and protein-PA attraction. The thermodynamics of this interplay is captured by a statistical fluctuation model, which allows the estimation of both energies. Quantification of the significance of each polar amino acid in the polybasic provides detailed insights into the molecular mechanism of the protein-membrane binding of LKB1. These results can be likely transferred to other proteins, which interact by intrinsically disordered polybasic regions with anionic membranes.

## Introduction

Protein-membrane binding, which is essential for many biological processes, takes place through a large number of transmembrane and peripheral proteins, which contain disordered regions in their structure.^1,2^ Some of these proteins are enriched in positively-charged residues or specific motifs, through which they are targeted to anionic membranes.^3–7^ These protein-membrane associations have been termed “fuzzy” since the protein remains unstructured during the binding process. These types of associations are naturally difficult to characterize and have been studied rarely. For instance, the extreme fuzzy association of the N-terminal region of ChiZ protein and its “specific” and “semispecific” binding to a POPG/POPE membrane have been described.^8^ The association of Src family kinases through the polybasic region in the disordered N-terminal domain has been also studied and the atomically detailed structural ensemble of the bound protein has been characterized.^9^ The importance of charged lipids for the interaction of the disordered RIT1 C-terminus with membrane has been also investigated.^10^ Although many important insights can be obtained from these studies, a detailed description of the structural properties and dynamics of these interactions are still missing, which are essential for a deep mechanistic understanding of the functional process of the association of disordered proteins with membranes.

Characterizing the fuzzy association of intrinsically disordered proteins (IDPs) is a challenging task. Experimentally, NMR spectroscopy can provide very precise information^8^ and MD simulations have become considerably more accurate for modeling the membrane association of the IDPs. However, despite the considerable progress in computational resources and force fields, the protein-lipid interaction studies remain elusive. Indeed, apart from the accuracy of the force fields there is a challenge of conformational sampling, particularly for IDPs, making these types of associations difficult to characterize.

Here we study the fuzzy association of LKB1, which is a serine/threonine kinase, with anionic membranes. LKB1 is localized to the plasma membrane (PM) by a disordered C-terminal phospholipid binding domain and its farnesylation motif. The farnesylation has been, however, reported not to be essential for the tumour suppression function of LKB1 in cultured mammalian cells or mice and Drosophila in vivo,.^11–13^ In contrast, the polybasic motif adjacent to the farnesylation motif is crucial for stable membrane recruitment and fully activation of the kinase activity of LKB1.^13^ The C-terminal polybasic region of LKB1 (aa 539–551), which includes eight charged amino acids(lysines and arginines (Ks,Rs)), facilitates a direct binding to PA and to much less extent to PtdIns(3,4,5)P3 and PtdIns(4,5)P2 lipids. ^13^ The mutation of these residues to alanine has been shown to abolish the protein-membrane binding in vitro as well as in cultured cells and in Drosophila in vivo. ^13^ The amino acid sequences for the wild-type (WT) and the mutated (MT) proteins with highlighted polar (in cyan) and mutated (in red) residues are shown in Table 1. However, cell biological studies also leave important questions unanswered, in particular, how exactly the binding of distinct amino acids within the polybasic motif reinforces binding of the entire protein to the membrane, how dynamic the protein-lipid binding is, and whether binding of LKB1 induces a local clustering of PA in microdomains, which in turn may strengthen the binding of the polybasic motif to the membrane.

**Table 1:**
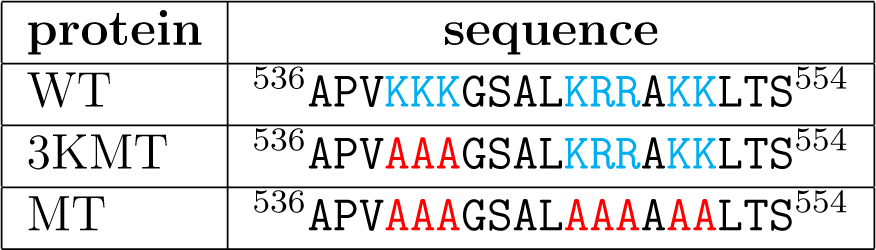
Sequences of the WT and MT proteins. The polar (cyan) and mutated (red) residues are highlighted.

Therefore, in this work we first probe the significance of PAs as well as single amino acids within the disordered polybasic region of LKB1 (aa 539–551) of the Drosophila LKB1 protein as well as its mutants with membranes containing various amounts of PA. For this purpose, we use extensive microsecond scale molecular dynamics (MD) simulations and multiple samples, alleviating the problem of sampling of the conformational space. The simulations are complemented by experiments in transfected cells with GFP-tagged variants of LKB1 and our systematic analysis shows the significance of PA for the protein-membrane binding and the effect of partial and whole mutation of the polybasic region.

Next, we study in more detail the fuzzy association of the polybasic region of LKB1 protein with anionic membranes and characterize the structural and dynamical properties of the protein-membrane binding. Using the spatio-temporal resolution of MD simulations, we specifically show the temporal changes in the structure of the protein as a result of the interaction with the membrane during the binding process and the stationary state at long time scale. We additionally discuss how the lateral distribution of PA molecules is affected by the presence of the interacting protein. We specifically describe the fluctuation of the dynamics of PAs using a theoretical model to understand the thermodynamics of the protein-membrane interactions from the perspective of the PAs. These results shed light on the mechanism of the protein-membrane interaction of LKB1 and the family of proteins interacting with membranes via polybasic regions, and in general, the interaction of IDPs with membranes including anionic lipids.

## Materials and methods

### System setup

For all simulations, we took a protein of 19aa from aa 536-554 of Drosophila LKB1 containing the polybasic motif in the C-terminus of the LKB1 protein. ^13^ The initial atomistic 3D structure of this protein segment was constructed using the SWISS-MODEL homology modeling web server.^14–16^ Accordingly, the model was built based on the 50S ribosomal protein L32 (PDB:1OND), with a sequence similarity of 37% and a sequence coverage of 66%. Note that the structure is supposed to be a disordered protein. Then CHARMM-GUI solution and membrane builder^17^ was used to prepare the systems with solvated proteins or the systems including also a membrane. During the process of the modeling, the protein is amidated and acetylated and finally solvated by water molecules with the addition of neutralizing ions, depending on the membrane composition and amount of the charge in the system. Concerning the salt concentration, we should mention that the binding of monovalent ions to the lipid headgroups is especially problematic and their binding affinity is not captured correctly by any of the force fields and their interaction with membrane is overestimated;^18,19^ therefore, we decided to only use the neutralizing ions. In the next step, the WT protein was simulated in solution in a few independent simulations for 200 ns. Then the structures at the end of these simulations were extracted and used to be combined with membranes of different compositions using CHARMM-GUI membrane builder. ^17^ The total number of lipids in the simulated systems is around 200. Additionally, two mutated models of the protein were constructed during the process of combining it with membrane: 1) the first three lysine residues are mutated to alanine (3KMT) and 2) all the lysine and arginine residues are mutated to alanine (MT). The amino acid sequences of the WT and the MT proteins are indicated in Table 1 and the simulated systems, including the lipid composition, number of samples and the simulation time for each sample are represented in Table 2.

**Table 2:** Simulation systems along with the protein type and membrane compositions as well as the number of samples and the simulation time are shown.

| system | protein | membrane | No. of PAs | No. of samples | sim time [μs] |
| --- | --- | --- | --- | --- | --- |
| WT-0%PA | WT | POPC100% | 0 | 7 | 2 |
| WT-5%PA | WT | POPC95%-POPS5% | 10 | 10 | 2 |
| WT-10%PA | WT | POPC90%-POPS10% | 20 | 13 | 2 |
| WT-20%PA | WT | POPC80%-POPS20% | 40 | 7 | 2 |
| 3KMT-0%PA | 3KMT | POPC100% | 0 | 5 | 0.15 |
| 3KMT-10%PA | 3KMT | POPC90%-POPS10% | 20 | 5 | 2 |
| MT-10%PA | MT | POPC90%-POPS10% | 20 | 7 | 2-4 |

### Simulation protocol

The MD simulations were performed using Gromacs version 2019.6.^20,21^ The CHARMM36m force field,^22,23^ which is a modified version of the CHARMM36 force filed for IDPs,and the TIP3P water model^24^ were used to define the interactions. Periodic boundary conditions were applied in all directions. The long-range electrostatic interactions were considered using the particle mesh Ewald method, ^25^ using the cutoff distance of 1.2 nm and the compressibility of 4.5*×*10*^−^*^5^. To treat van der Waals (vdW) interactions, the cut-off schemes with the cutoff distance of 1.2 nm were used, which is smoothly truncated between 1.0 and 1.2 nm. The electrostatic interactions were considered using the particle mesh Ewald method. ^25^ The constant pressure for the protein-membrane systems was semi-isotropically maintained at 1 bar with the use of Berendsen^26^ and Parrinello-Rahman barostats,^27^ respectively for the equilibration and the production simulations. The constant temperature at 310 K was controlled by coupling the system to the Nośe-Hoover thermostat.^28,29^ The bonds were constrained using the LINCS algorithm.^30^ Before the equilibration process, all systems were first minimized in 10000 steps. The protein-membrane systems were subsequently equilibrated using initially the NVT (2ns) and then the NPT (28 ns) according to the input files provided by CHARMM-GUI. During the course of equilibration, restraints (starting with 4000 kJ/mol*^−^*^1^*·*nm*^−^*^2^) were applied on the heavy atoms of the protein and the lipids, which were in multiple step gradually decreased to 50 kJ/mol*^−^*^1^*·*nm*^−^*^2^. The production simulations were performed for 2 *µ*s using a time step of 2 fs. For the MT system the simulation were continued for 4 *µ*s. In order to provide statistically independent simulations, for each system different samples were run using different structures of the protein extracted from the simulations of the protein in a solution without the presence of membrane. For these simulations, the same protocol was used as the protein-membrane systems except that the isotropic pressure coupling was utilized and for each simulation the production simulations were performed for 200 ns.

The simulation results were analyzed using in-house python codes, incorporating the MDAnalysis package^31,32^ and the GROMACS tools. VMD was employed to visualize the trajectories and prepare the snapshots. ^33^

### Order parameter

The order parameter of the lipid chains was calculated according to ref.,^34^ using molecular order parameter formula defined as

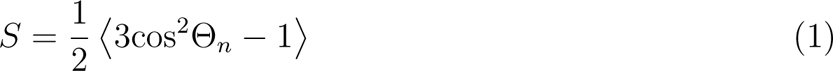

where Θ*_n_* is the angle between the vector constructed by n^th^ segment of the hydrocarbon chain, i.e. *C_n−_*_1_ and *C_n_*_+1_, connecting the *n −* 1 and *n* + 1 carbon atoms, and the membrane normal (z-axis). The angular brackets represent the time and ensemble average.

### Cell culture and transfection

Schneider S2R+ cells on coverslips were transfected with GFP-LKB1 variants under a ubiquitous promoter (Ubi::GFP-LKB1) using FUGENE (Promega).^35^ 48 h after transfection, cells were fixed with 4% paraformaldehyde in PBS pH7.4 for 20min. Subsequently, cells were washed three times with PBS and nuclei were labelled with DAPI for 20min. Cells were mounted in Mowiol and imaged with a confocal microscopy (Leica SP8).

## Results

### Results: Protein perspective

#### PA is essential for the protein-membrane binding of LKB1

In all systems the protein is first in solution at the beginning of the simulations, so that the distance of the center of mass (COM) of the protein from the center of the membrane along the membrane is 6 nm (Figure 1A top) and the protein can freely move. Depending on the membrane composition, the protein can be attracted to the membrane and interact with it (Figure 1A middle,bottom). In order to check if the protein interacts with the membrane we plotted the projected distance of the COM of the protein and the closest heavy atom of the protein from the center of the membrane along the membrane normal (z-axis) (Figure 1B-F). For membranes without PA lipids (WT-0%PA system, Figure 1B), the protein can occasionally approach the membrane and interact with it, but no persistent protein-membrane binding takes place. However, as soon as 5% PA lipids (or more) is added to the membrane persistent interactions of the protein with the membrane occur (Figure 1C). Thus, a small fraction of PA lipids is sufficient for a stable protein-membrane binding.

**Figure 1:**
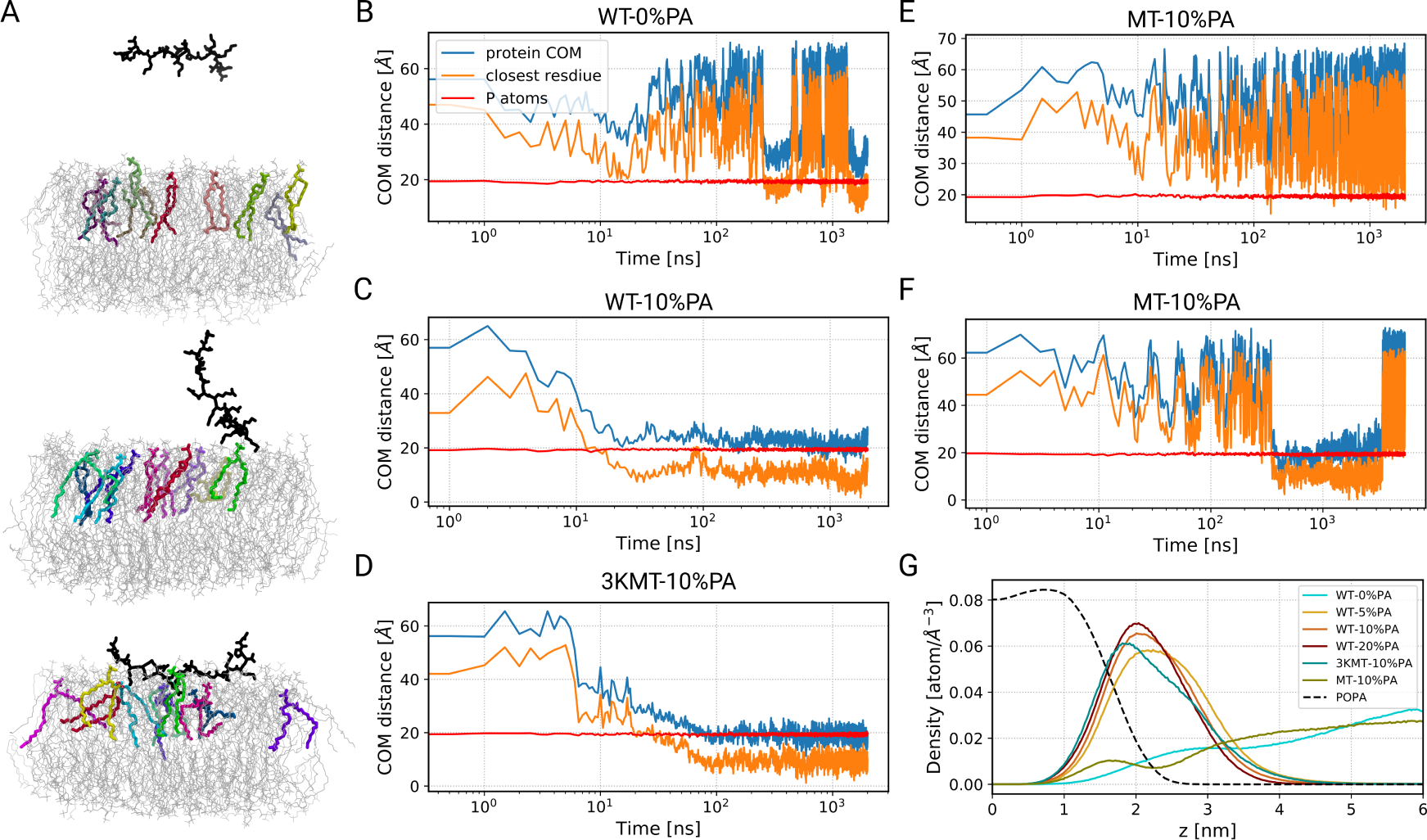
Presence of PA and mutation of the polybasic region affect the protein-membrane binding. A) The snapshots of the protein-membrane system for the WT-10%PA system at different times of the simulation are shown. The protein and PAs in one leaflet are shown respectively in black and colored licorice representation. The PC lipids are shown as grey lines. B-F) The distance of the COM of the protein with respect to the center of the membrane (blue), the closest heavy atom of the protein (orange) and the average position of the P atoms of the lipids (red) projected on the membrane normal (z-axis) as a function of time for different systems are shown. E and F correspond to two different realizations. G) The density of the protein in different systems as well as the density of PA lipids (dashed line) are represented.

Interestingly, the mutation of the first three lysine residues to alanine (3KMT-10%PA) does not abolish the stable interaction with the membrane (Figure 1D). Conversely, when all the lysine and arginine residues are mutated to alanine (MT-10%PA), either no interaction with the membrane takes place (Figure 1E) or only short binding periods (Figure 1F).

In order to confirm that our simulations represent the behavior of the protein in vivo, we transfected cells with GFP-tagged variants of LKB1. Indeed, while wild type full length LKB1 or a truncation of LKB1 containing the last 49aa (LKB1 518-C) accumulates mainly at the plasma membrane, mutation of all positively charged amino acids (LKB1ΔLB = Δlipid-binding) abolishes membrane binding resulting in a cytoplasmic accumulation of the mutant protein (Figure 2A, B, D). Strikingly, as predicted from our simulations, mutation of the first three lysines (3KMT) did not substantially affect cortical localization of LKB1 (Figure 2C).

**Figure 2:**
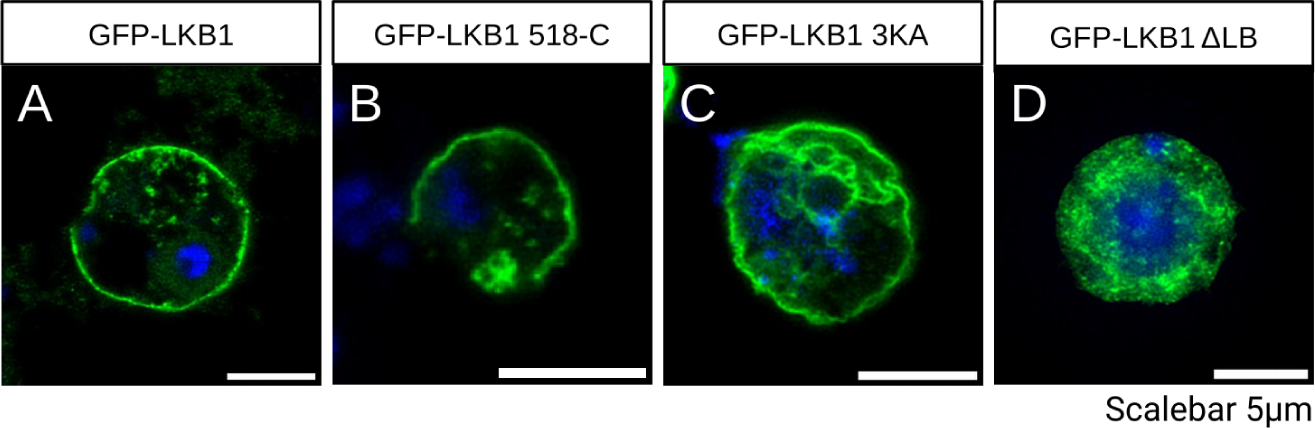
Mutation of the lipid-binding motif results in disturbed membrane association of LKB1. A-D) Schneider 2R+ (S2R+) cells were transfected with GFP-LKB1 variants and stained with DAPI. Wild type LKB1 (A) and the isolated C-terminus (amino acids 518-566, B) accumulate predominately at the plasma membrane, whereas mutations of all basic amino acids results in cytoplasmic aggregates (D). LKB1 with a mutation of only three basic amino acids within the polybasic motif is still robustly targeted to the cortex (C).

#### Protein’s embedding in membrane

The general protein-membrane interaction is also reflected in the density plots of the protein and lipids (Figure 1G), showing to what extent the protein can be attracted to the membrane. The density plots show that the protein with the higher amount of PA lipids in the membrane can systematically penetrate the membrane more efficiently. Interestingly, for the 3KMT-10%PA system, the protein can approach the membrane stronger than the WT one (WT-10%PA), while some parts of it remain further separated from the membrane due to the mutation. As expected, both MT-10%PA as well as WT-0%PA display even repulsive interaction with the membrane. The observation from Figure 1F about the temporary binding of the 3KMT-10%PA system is reflected by the small maximum around 1.5 nm.

#### Protein’s structural changes

Next we looked at how the structural properties of the protein is affected as a result of the interaction with the membrane. The root-mean-squared deviation (RMSD) in steps of 50 ns for the WT-10%PA system (Figure S1A) and the average of RMSD values for all systems (Figure S1B) are shown. The RMSD for WT systems dramatically decreases for pure membrane system and only slightly decreases for the WT systems with increasing amount of PA. The 3KMT-10%PA system shows the highest RMSD value, which is due to the presence of the weakly bound mutant part of the polybasic motif that causes the protein to be highly flexible.

Since the protein does not have a well-defined secondary structure, its structure dramatically changes as a result of the protein-membrane interaction. Therefore, we analysed these structural changes in more detail: The average of the sum of squared distances of CA atoms of the protein amino acids from the COM of the protein backbone, once projected along the z-axis (Δz) and once on the xy-plane (Δr), i.e. the respective standard deviations was calculated. These values were then normalized by the corresponding values for the free proteins inside the solution in the systems without PAs (WT-0%PA and 3KMT-0%PA), which do not interact with the membrane (Δz*_norm_* and Δr*_norm_*). Therefore, a value of unity represents a property close to that of the free protein in solution without the interaction with the membrane. As the protein binds to the membrane, Δz*_norm_* gradually reduces and adopts values below 1. This reflects a generic behavior that along with the binding processes of amino acids with the membrane a flattening of the protein takes place. Interestingly, in the long-time range the normalized standard deviation increases again (Figure 3A). A slightly different behavior is observed for the 3KMT-10%PA system. Here the time-averaged normalized standard deviation is close to unity. This may be related to the fact that the mutations reduce the strength of the interaction with the membrane. Along with this interpretation the reduction in the standard deviation Δz is most pronounced for systems with a higher concentration of PAs (Figure 3A right).

**Figure 3:**
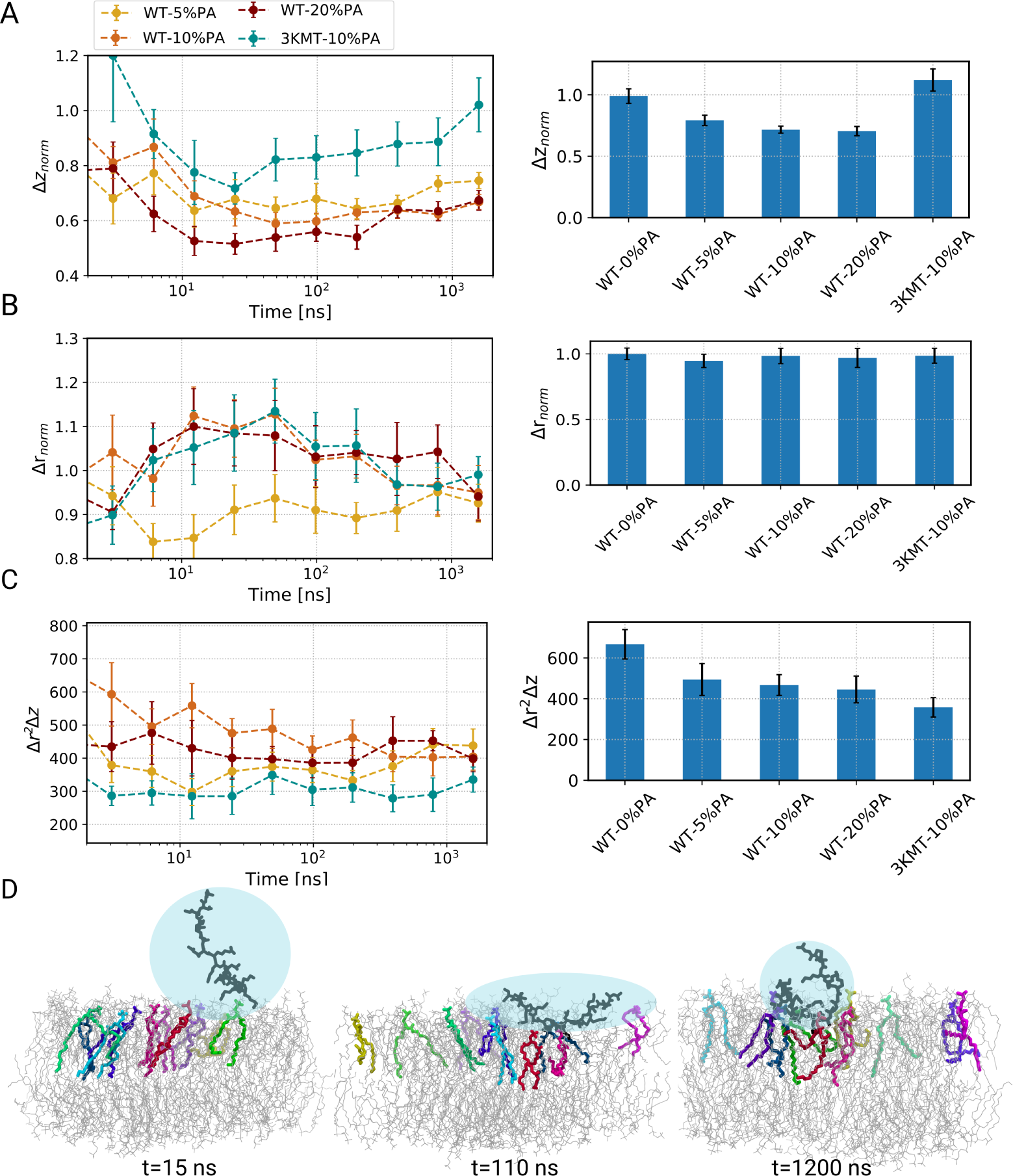
Structural properties of the protein changes during the course of membrane binding. A) The root mean squared of the average distances of CA atoms of the protein from the COM of the backbone atoms of the protein projected on the z-axis (Δz*_norm_*) and B) the xy-plane (Δr*_norm_*). C) The volume of the protein, defined as Δr^2^Δz, i.e., without any normalization, are shown over blocks of the simulation time. The corresponding average values of the quantities in A-C for the last 500 ns of the trajectory are represented in bar plots. D) The snapshots of the protein on the membrane, showing various representative structures, at different times of the simulation for the WT-10%PA. The protein is shown in black and the PAs in the interacting leaflet in different colors, using licorice representation.

For all systems except for WT-5%PA the value of Δr*_norm_* conversely increases as a result of the interaction for intermediate times and for longer times approaches the value, seen for the non-interacting case (Figure 3B). To a good approximation this behavior is opposite to the time dependence of Δz*_norm_*. This stretching increases the free energy of the protein and is likely counterbalanced by the enthalpic interaction with the membrane in particular with PA. The disappearance of this stretching effect for long time periods (Figure 3B right) suggests that some reorganization of the PA distribution occurs, which we will address further below. The opposite trend for the time-dependence for WT-5%PA may be related to the lower number of PAs.

Next, we analyze Δr^2^Δz, reflecting the effective volume of the protein (Figure 3C). While the time-dependence is somewhat difficult to interpret, we clearly see that for the WT the final volume is quite insensitive to the PA content. However, as compared to the WT without significant binding effects, i.e. with no PA, the volume shrinks by approx. 25%. The different volume for 3KMT-10%PA reflects the different sequence of this protein.

Finally, in Figure 3D we show the time-dependence of the conformation for the representative example of WT-10%PA. While for short times no direction is preferred, for intermediate times one can see that the protein adopts the shape of an oblate spheriod. This is compatible with the increase of Δr^2^ and the decrease of Δz. In the equilibrated states the shape becomes somewhat more spherical but with a smaller volume compared to the initial unbound state.

### Results: Protein - PA interplay

#### The polybasic motif of LKB1 mainly binds to PAs

The structural changes in the protein structure, discussed in the previous section, indicate the presence of a subtle interplay of protein conformation and PA arrangement. Therefore, in this section we analyzed the protein-membrane interaction in more detail, starting with the percentage of lipids interacting with the protein throughout the simulations. The number of interacting lipids was based on the number of contacts using a cut-off distance of 4.0 ^°^*A* between any atoms of the protein and the lipids headgroup. The results show that the protein binds to the membrane in the first few nanoseconds. Interestingly, when the amount of PA is lower than or equal to 10%, the number of lipids reaches a final value quite fast (Figure 4A,B), whereas in the case of WT-20%PA, it takes a comparatively long time until an optimum interaction state is achieved (Figure 4C).

**Figure 4:**
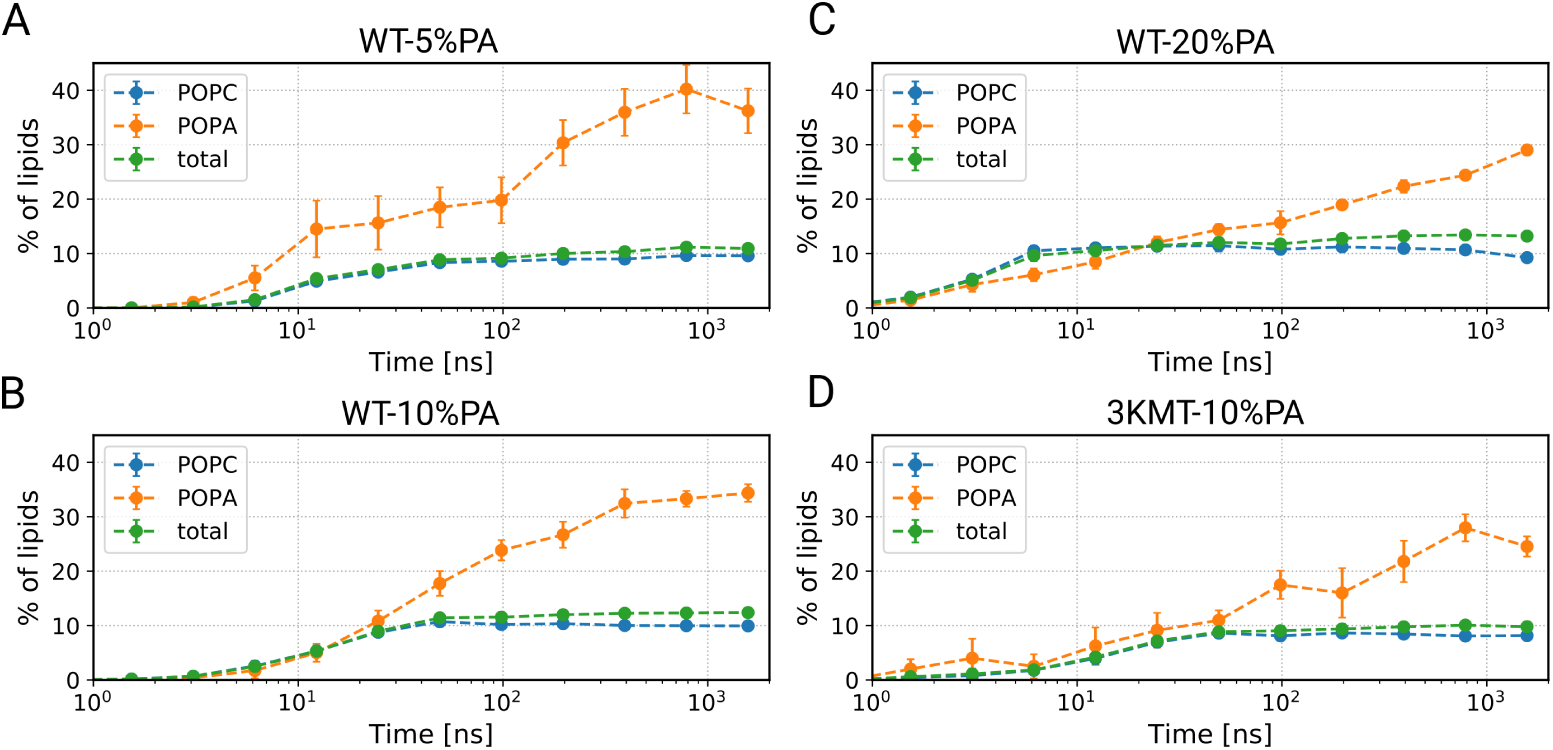
Time evolution of the percentage of interacting PA with the protein. A-D) The average of the percentage of different lipid types interacting with the protein for different systems is shown.

If there were well-defined binding sites for PAs within the polybasic motif, one would expect that the absolute number of PA molecules close to the protein should be insensitive to the PA concentration. Then the fraction of bound PAs would correspondingly decrease for increasing PA concentration. However, we see that there is only a very small decrease of the fraction of bound PA (approx. 35% for WT-5%PA and slightly smaller values for WT-10%PA and WT-20%PA). This implies a strong increase of the absolute number of PAs with increasing PA concentration. As a consequence, the attraction of the PAs may take longer for higher PA content. As expected, the 3KMT protein is not able to interact with the same number of lipids as in the WT system due to the point mutations (Figure 4D).

To explore the translocation process in more detail, we calculated the 2-dimensional mean-squared displacement (MSD) of the protein and PAs in different systems along the membrane plane for the last 500 ns of the simulations. The MSD plots of the COM of the alpha carbon atoms of the protein show that in the WT-0%PA system, where the protein does not interact with the membrane, the protein diffuses almost an order of magnitude faster (Figure S2). The mobility of the proteins, which interact with the membrane is similar for all systems and is very close to the mobility of the PAs and show a diffusive behavior (Figure S2). This observation indicates a strong dynamic correlation of protein COM and PA dynamics.

#### Importance of amino acids in protein-membrane binding

Next we studied the protein-membrane binding on the level of single amino acids in the polybasic region of LKB1. To characterize these properties in the long-time stationary regime, we averaged the fraction of bound PAs during the last 500 ns of the simulation (Figure 5A-D). One can clearly identify three groups of positively charged residues of the protein, i.e., KKK, KRR and KK, which display a particularly strong attraction of PAs. The respective time-dependencies (summed over each group) are additionally displayed in Figure S3. As already known, during the simulation time no complete convergence of the WT-20%PA binding is reached so that actual number of contacts in the long-time limit is expected to be higher.

**Figure 5:**
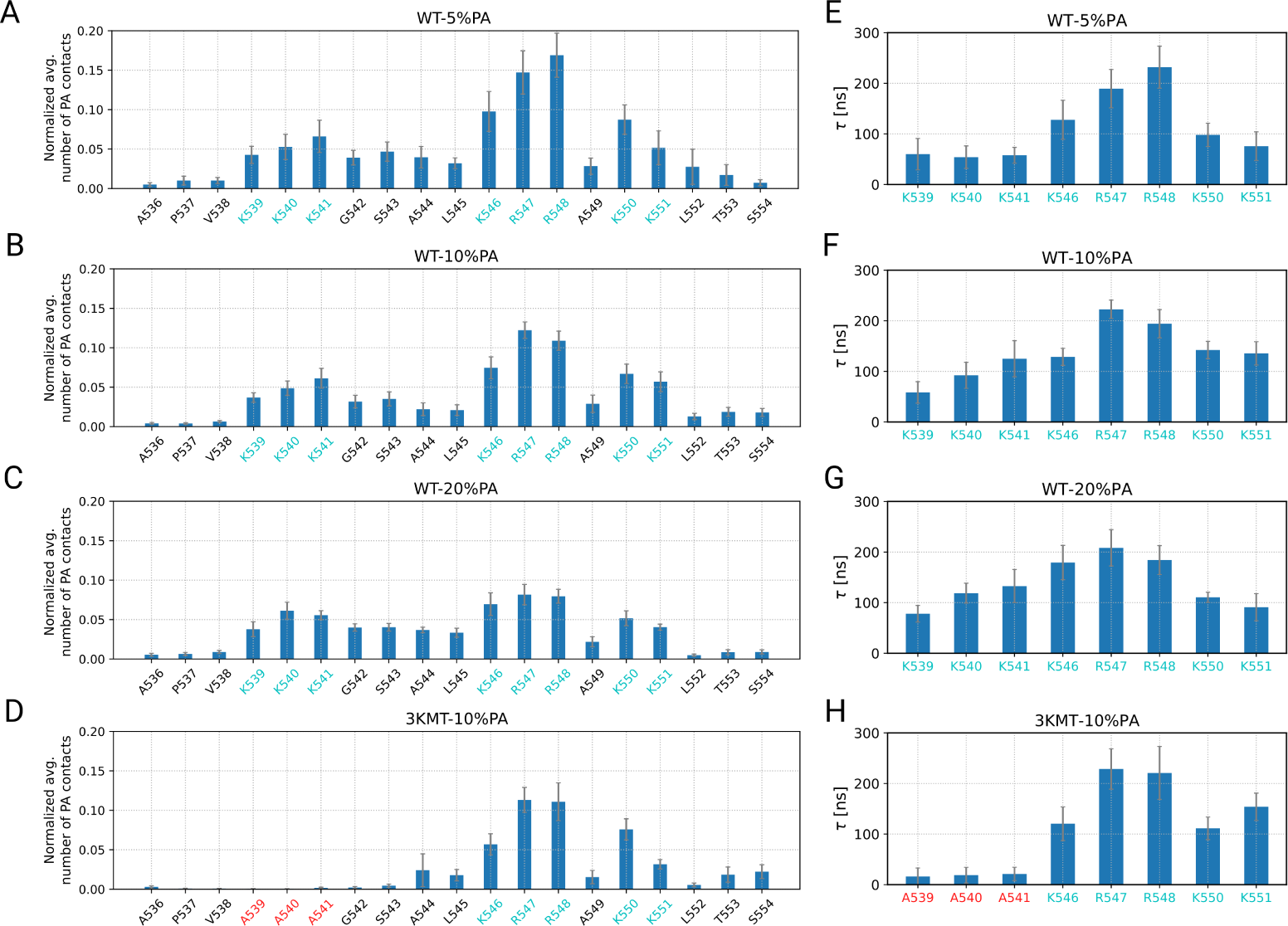
The average number of contacts of PAs with the protein residues is in agreement with the residence time of PAs. A-D) The average number of the PA contacts with the protein residues normalized by the number of PAs in the interacting leaflet is shown over the last 500 ns of the simulation. E-H) The average residence time of PAs with the amino acids in the polybasic region of the protein. The residence times were calculated from the auto-correlation function of the time series of the number of contacts of PAs with the polar residues.

The KRR group establishes the highest number of contacts with the PAs, which is likely due to the presence of arginine residues in this group. The other two groups show a somewhat smaller but still significant affinity to the membrane. As already discussed in the context of Figure 4, the fraction of bound PAs does not change much despite the enormous (factor of 4) variation of the PA concentration. Only for the KRR group the fraction of lipid contacts somewhat increases for smaller PA content. This suggests that there are specific PA-binding sites in this group.

As expected, the 3KMT-10%PA system shows a negligible number of contacts of the mutated residues with the PAs whereas the other groups show a similar behavior as the wildtype.

Moreover, it is also interesting to probe with how many PA lipids each polar residue interacts (Figure S4). The results obviously depend on the amount of PAs in the system. For the WT-5%PA system, for instance, at most two PAs can bind to these residues (Figure S4A), whereas for the WT-10%PA system the arginine residues and K550 can bind also to 3 PA lipids at the same time (Figure S4B). For the WT-10%PA system, almost all the residues can bind to three PAs (Figure S4C), while for the 3KMT-10% system this is mainly the case only for the arginine residues (Figure S4D).

Additional information can be gained from the analysis of the residence times of an individual PA molecule at specific binding site, based on the auto-correlation function of the time series of the protein-lipid contacts (Figure 5E-H). Interestingly, in agreement with the structural properties, the second group of amino acids in the polybasic region, i.e., KRR, shows longer residence times, which is again likely due to the presence of arginine residues in this group. This highlights the expected correlation between high binding affinity and long residence times. For comparison we also calculated the residence time of PAs with the entire protein, which is around 350 ns, and thus, approximately only three times longer than the average residence time with the individual residues. This shows that there is a gradual exchange of the PAs, which interacts with the protein, highlighting a highly dynamic binding. At the same time it is interesting to compare the MSD at 350 ns (which is approximately 60 nm^2^) (Figure S5) with the projected area of the protein on the membrane (which is approximately 93.6 nm^2^ for the case of the WT-10%PA system). These values are relatively close to each other, which implies that a scenario where the bound PA diffuse together with the protein over distances much longer than the protein size can be excluded (Figure S6A). Instead, the protein-lipid binding has a restricted temporal-space behavior (Figure S6B).

### Results: PA perspective

#### Agglomeration of PAs

Since the polybasic motif of LKB1 interacts mainly with PAs, this suggests that the protein can induce the agglomeration of PAs and as a result affect the lateral distribution of PAs. To probe this effect in more detail, we first calculated the radial distribution function (RDF or *g*(*r*)) of PAs along the xy-plane (Figure 6). To compare the effect of the protein-membrane interaction, we considered the MT-10%PA system as a control system since the protein does not interact with the membrane and no change in the behavior of PA should be observed. We always used the last 500 ns of the simulation to be sensitive to the stationary situation. Without the impact of the protein the PAss are strongly repelling each other for both leaflets (Figure 6A). This is a consequence of the mutual Coulomb repulsion. Remarkably, it gives rise to a radial distribution function *g*(*r*) *<* 1 for basically all distances. In contrast, in the presence of bound LKB1 (Figure 6B-D) one observes some agglomeration effects, i.e. *g*(*r*) *>* 1, in the upper leaflet (interacting leaflet), whereas the PA structure in the lower leaflet is basically unmodified. This is due to the fact that the protein tends to interact with PAs and, as a consequence, it induces their agglomeration. A typical realization can be found in Figure 6E.

**Figure 6:**
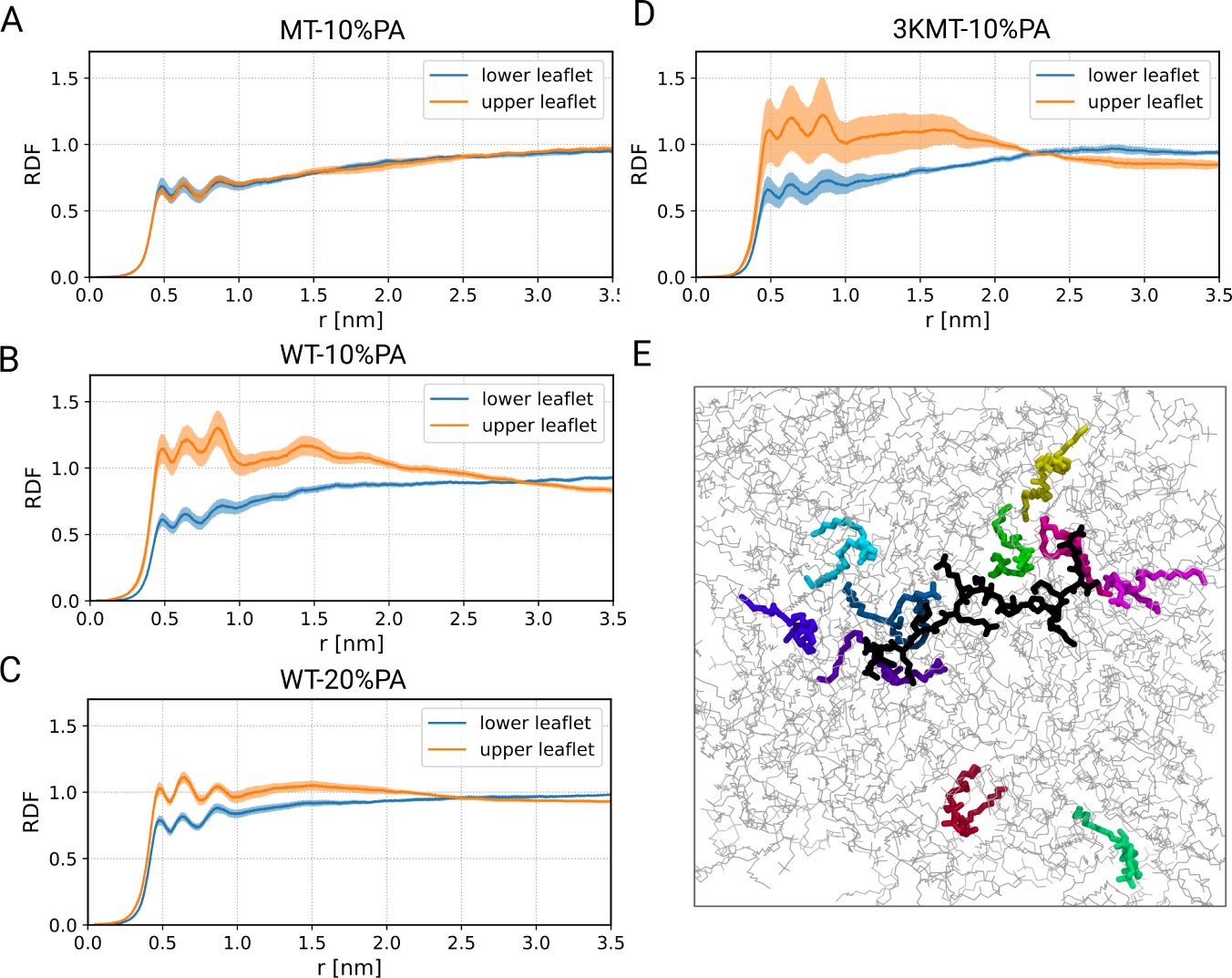
Protein-membrane interaction results in accumulation of PAs. (A-D) The RDF profile of PAs in the upper leaflet (the interacting leaflet) and the lower leaflet for the last 500 ns of the simulation for different systems. E) A snapshot of the protein-membrane system (WT-10%PA) from the top view is shown. The protein and the PAs in the interacting leaflet are shown respectively in black and colored licorice representation. The PC lipids are shown in grey lines.

Furthermore, in Figure S7 we plot the time evolution of the first peak of the RDF. As expected, the observed increase with time reflects the increase of binding PAs, as shown in Figure 4. Interestingly, for the MT-10%PA system the peak height decreases with time. Thus, it takes time to accomplish a configuration where the PAs efficiently repel themselves

Next, we studied the number of PA molecules interacting with the polybasic motif of LKB1 for the cases of the WT and 3KMT-10%PA systems. From the last 500 ns of the simulation we estimated their average number *µ_sim_* as well as their fluctuations, expressed by the standard deviation *σ_sim_* of their distribution. To find an interpretation of the size of the fluctuations, we compared these values with the expectation when assuming a binomial distribution, i.e. 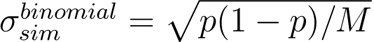, where *p* is the probability that PAs interact with the protein and *M* is the number of PAs. The values for *µ_sim_* and 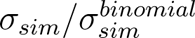 are listed in Table 3. Interestingly, the actual fluctuations are somewhat smaller than the predictions of the binomial distribution. This effect is particularly pronounced for membranes with a higher PA concentration.

**Table 3:** Fluctuations of interacting PAs: Comparison of simulations with the predictions of the fluctuation model.

| **system** | **$r$** | **$\mu_{sim}$** | **$\sigma_{sim}/\sigma_{sim}^{binomial}$** | **$\mu_{model}$** | **$\sigma_{model}/\sigma_{model}^{binomial}$** |
| --- | --- | --- | --- | --- | --- |
| WT-5%PA | 7.2 | 38.5 | 0.90 | 38.5 | 0.90 |
| WT-10%PA | 7.4 | 34.5 | 0.81 | 32.0 | 0.82 |
| WT-20%PA | 7.7 | 28.6 | 0.64 | 24.4 | 0.72 |
| 3KMT-10%PA | 6.4 | 26.3 | 0.78 | 29.1 | 0.78 |

To obtain a mechanistic understanding of the protein-membrane binding, we formulated a minimal model, incorporating the following key effects. (1) Without binding and repulsion, the distribution is a binomial distribution. The probability *r* of a lipid to be bound can be estimated from the MD simulations when analysing whether a randomly placed lipid is counted as a bound or an unbound lipid. The values for *r* slightly depend on the chosen protein and PA composition and are listed in Table 3. (2) A binding PA molecule experiences a binding energy *E*_1_ (here expressed relative to temperature). (3) There exists a maximal number *n_max_* of lipids. For reasons of simplicity this value is chosen as the number of polar amino acids in the polybasic region, i.e. *n_max_* = 8 for the WT and *n_max_* = 5 for 3KMT. (4) Bound lipids repel each other due to Coulomb interaction. If at a given time one has *n* bound PAs, this can be expressed (again relative to temperature) as *E*_2_(*n −* 1)^2^ where the quadratic dependence corresponds to a mean-field approximation. We denote the resulting model as *fluctuation model*.

The probability to have *n* bound PAs is proportional to 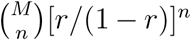 exp[*E*_1_ *· n − E*_2_ *·* (*n −* 1)^2^] if *n ≤ n_max_* and 0 otherwise. From this distribution the different moments of the distribution of bound lipids can be easily calculated. The two adjustable parameters *E*_1_ and *E*_2_ can be determined from reconstructing the first and second moment for WT-5%PA, yielding *E*_1_ = 2.33 and *E*_2_ = 0.12. Indeed, for *E*_2_ = 0 there are no deviations from the binomial distribution because the upper limit *n_max_* = 8 does not matter for *M* = 5. Thus, both the repulsion of lipids as well as the presence of a maximum number of bound lipids reduces the fluctuations as compared to the binomial distribution. For the estimation of *E*_1_ one can obtain an analytical approximation from the relation *r/*(1 *− r*) exp(*E*_1_) = *µ/*(1 *− µ*), which strictly holds for *E*_2_ = 0. For WT-5%PA this yields *E*_1_ = 2.08 which is close to the actual fitted value of 2.33.

The predictions of the fluctuation model for the other three cases are shown in Table 3 and the two adjustable energy parameters, representing attraction and repulsion effects, as obtained from the WT-5%PA system. One can indeed find a very good agreement between the actual numerical data and these predictions for all three cases. Thus, one may conclude that to a reasonable approximation the binding properties can be related to this simple fluctuation model. Two key conclusions can be drawn. First, the model analysis shows that both due to the stronger repulsion effects upon crowding as well as due to the presence of a maximum number of binding sites, the fluctuations are reduced as compared to the binomial distribution, as also seen for the simulation data., indicating the importance of these effects. Second, the description of the observed fluctuations of a relatively small system via the fluctuation model allows for a straightforward estimation of the relevant energy parameters, expressing the typical binding (free) energy as well as the repulsive energy.

The agglomeration of PAs can be additionally observed in the density maps of PAs, considering the protein at the center of the density map and looking at the position of lipids around it (Figure 7A). It is clearly observed that for the lower leaflet the lipids are distributed homogeneously (Figure 7A right), whereas for the upper leaflet a considerable accumulation of PA is obvious (Figure 7A left).

**Figure 7:**
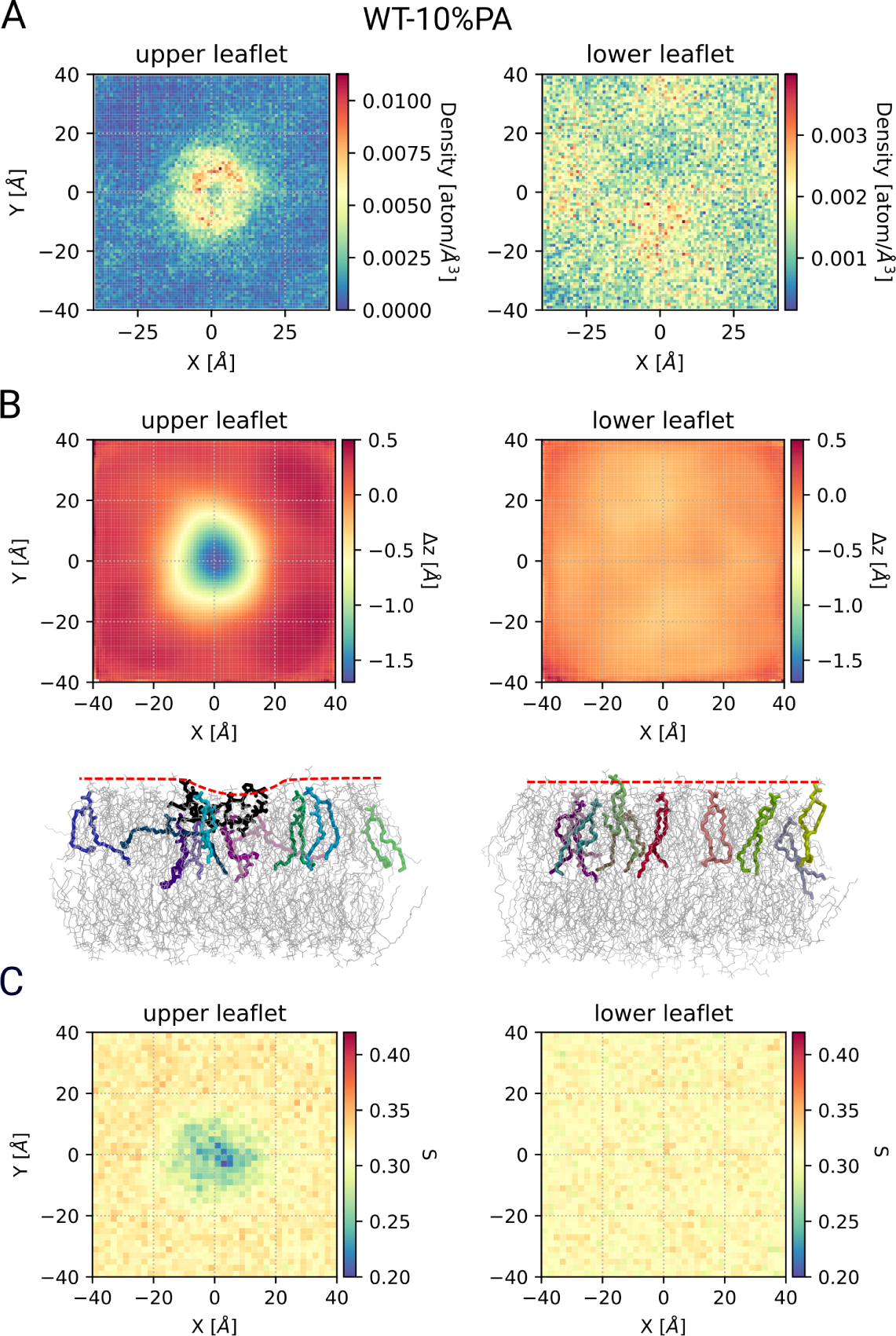
Density of PA and membrane properties. A) Density maps of PAs for the WT-10%PA system when considering the protein at the center of the membrane. Therefore, the density maps were calculated by changing the coordinates in a way that the protein is kept always at the center of the membrane along the xy-plane. B) The height profiles of the upper and lower leaflets for the WT-10%PA system, when considering the protein always at the center of the membrane, are shown. Only the position of P atoms of PC lipids was used for these calculations. Subsequently, for each leaflet the average COM of the P atoms in the respective leaflet was subtracted from the height positions along the membrane normal. The snapshots of the protein-membrane system for the interacting (left) and non-interacting (right) are shown. The approximate surface of the membrane is shown by a red dashed line. The protein and PA molecules in one leaflet are shown respectively in black and colored licorice representation. The PC lipids are shown as grey lines. C) The order parameter of PA and PC lipid chains for the WT-10%PA system again when considering the protein at the center of the membrane.

The interaction of the polybasic motif of LKB1 with the membrane can also affect the membrane properties. Here we study the height profiles of the lipids’ headgroups and the resulting thickness of the membrane. We calculated the height profiles again by placing the protein at the center of the membrane and calculating the height profiles of the P atoms of the PC lipids (Figure 7B). Interestingly, we see that the surface of the membrane close to the position of the protein is around 2 ^°^*A* deeper than the other parts, meaning that the protein is somehow absorbed by the membrane. This is likely associated with additional structural changes in the lipids structure, i.e., the order parameter of the lipid chains, which adopts lower values for the lipids in the vicinity of the protein (Figure 7C).

## Discussion

In this work, we probed the interaction of the C-terminus of LKB1, which adopts a disordered structure with membranes containing different concentrations of PA. PA turns out to be imperative for the protein-membrane binding. This is in agreement with cell culture experiments using GFP-LKB1 variants. These observations are similar to the RIT1 protein, which is also localized close to the PM and lacks the C-terminal prenylation and helps many other subfamily members to adhere to cellular membranes. The MD simulations of the disordered C-terminus of tRIT1 revealed a dependency of the membrane interactions on the lipid composition in membranes containing POPC/POPS.^10^ We additionally showed that mutation of the first three lysines within the polybasic motif (3KMT) did not prevent the protein-membrane binding, consistently seen from MD simulations and cell culture experiments. Furthermore, we showed that the middle group of residues in the polybasic region plays a more important role than the adjacent groups, which might be associated to the presence of arginine residues in this group, as this amino acid has been shown to strongly attract phosphate and establish extensive H-bonding.^36^ Consistently, higher residence times of PA molecules were observed with arginine residues compared to lysine residues.

We would like to stress that for simulations of IDPs it is quite some challenge to achieve reasonable agreement with experimental data.^23,37–42^ To achieve sufficient configurational sampling we therefore performed simulations in the microsecond regime and used multiple samples in order to reach and also characterize the stationary state after the binding process. Furthermore, we profited from the recent improvement of the force fields for IDPs,^23,40,41,43^ particularly due to the dispersion-corrected water models,^44^ which have resulted in better solvation, and therefore, more realistic simulations.

Recently, Hicks et al. have characterized the membrane association of the disordered, cytoplasmic N-terminal region of ChiZ, an Mtb divisome protein, by combining solution and solid-state NMR spectroscopy with MD simulation.^8^ They suggest the hydrogen bonding between arginine residues and POPG lipids as a driving mechanism for membrane association. The general interaction is called “semispecific” since there is no specific binding interface and the protein did not take any secondary structure upon membrane binding. This goes along with a highly dynamic protein-membrane binding scenario. Furthermore, the authors argue that this driving mechanism is part of the nonrandom characteristic of the fuzzy interaction. In contrary, the *α*-synuclein disordered proteins and the N-terminal region of M. tuberculosis FtsQ partly form *α*-helices when bounded to membranes.^2,45^ Likewise in our case the arginine residues locate in the middle of sequence and take a major role in the protein association. Furthermore, we also find this very dynamic behavior.

We believe that some directions of the current work are of particular general relevance:

### Initial binding process

Characteristic behavior emerges during initial binding process of the protein until a stationary state is reached. The compactness of the protein along the membrane normal and the membrane plane is modified throughout the binding-process in a non-monotonous fashion. In initial period the protein acquires an oblate form. In this way, an efficient search for an optimal interaction with the PA of the membrane is possible. Afterwards the flatness becomes reduced, going along with the clustering of PA lipids. Thus, one can clearly find a non-monotonous behavior until a stationary state is reached. During the whole process the effective volume of the protein is continuously decreased. All these observations are made possible due to the large conformational entropy of IDPs.

### Fuzziness of the stationary binding

From analysis of the spatio-temporal fluctuations of the protein *and* of the acid lipids we observed that during the time of a typical PA-protein binding process the protein moves at most a distance close to its size. Due to the protein-membrane interaction and its dynamical nature, the arrangement of PA molecules in the membrane are changed and showed a dynamic clustering in the proximity of the protein. This agglomeration was restricted to the interacting leaflet.

### Binding strength

The binding strength of a protein with a well-defined structure with a membrane is usually determined via PMF calculations.^46–48^ Interestingly, in the work of Yamamoto et al. it is observed that the binding energy decreases with the increase in the number of acidic lipids in the membrane; it is, however, not clear how many lipids actually interact with the protein in each case.^49^ Remarkably, in the work of Larsen et al. extensive FEP calculations have been performed to extract the free energy of binding also from the perspective of the acidic lipids. For their example of PIP2, the authors obtained the binding free energies between 1.6 to 8.5 *k_B_T*, depending on the binding site of the interacting protein and the protein itself.^46^ Summing over the different binding sites yields free energy binding energies which are compatible with the binding free energies of the respective protein. Furthermore, significantly stronger binding is expected for PIP2 as compared to PS.^46^ For our disordered protein we refrained from the PMF calculations due to the sampling problem of the large configurational space. Therefore, here we introduced a much simpler method to extract information about the binding free energies. Starting from the lipids perspective we analysed the fluctuations of the number of interacting PAs and from this we derived an estimation of the binding free energy for an individual PA, resulting in a value of 2.3 *k_B_T*. From our fluctuation analysis we could even estimate the effect of reduced binding with the increase in the number of lipids, in agreement with the observations in the work of Yamamoto et al.^49^ This effect is likely related to the Coulomb repulsion among the different PAs.

### Membrane properties upon binding

In general one expects that the protein-membrane interaction and the resulting domain formation in the membrane upon adsorption of the protein affects the membrane properties. Indeed, clustering of anionic lipids and the analogous change in membrane properties was observed in the MD simulations of strong polycations with a POPC/POPS membrane^50^ and for the case of polyethylenimines with a membrane, containing POPC/DOPA.^51^ Here we could specifically reveal the decrease in thickness at the location of the protein and the decrease of the order parameter of the lipids’ acyl chains. Thus, these observations are not only characteristic for synthetic polycations but also for IDPs.

## Conclusions

In summary, we revealed that the polybasic motif of LKB1 can establish stable, albeit highly dynamic, binding with membranes with low amounts of PA. This suggests that farnesylation of LKB1 is not specifically required for stable association of LKB1 with membranes, which has been supported by in vivo experiments (Dogliotti et al.^13,35^). Furthermore, the LKB1-membrane interaction revealed interesting mutual effects of the protein and anionic lipids, such as proteins’ structural modifications and agglomeration of PAs, providing microscopic and thermodynamic insights into the interaction of LKB1 with membranes containing PAs. Due to the generality of our observations, we expect that these insights also help to understand the interaction of other IDPs, containing polybasic regions, with membranes, containing anionic lipids.

## Acknowledgement

The authors thank the high performance computer resources at University of Muenster (Palma) and the financial support by the German Science Foundation (DFG, projects CRC1348-A01 and CRC1348-A05) and the Interdisciplinary Centre for Clinical Research (IZKF) Münster (Kr-A-031.21).

## Supporting Information Available

The following files are available free of charge.

Figure S1. RMSD of the protein backbone in slices of 50ns. Figure S2. MSD of protein and PAs. Figure S3. Interaction of PA with different blocks of residues in the polybasic region. Figure S4. Probability of the average number of PAs interacting with polar residues of the protein. Figure S5. Residence time of PA with the protein. Figure S6. Two scenarios for the protein-lipid binding. Figure S7. The average heights of RDF profiles.

## TOC Graphic

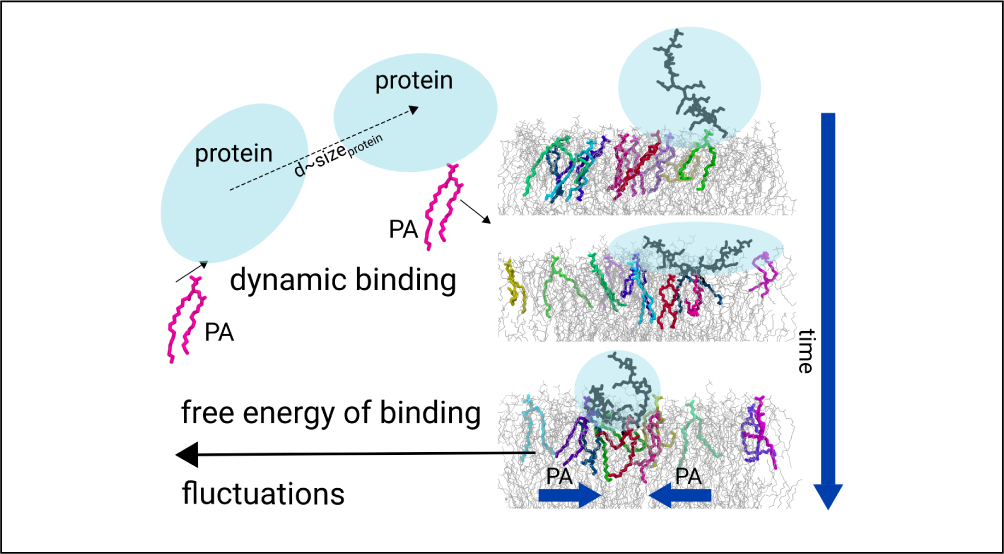

