## Supporting Information for "Elucidating the Membrane Binding Process of a Disordered Protein: Dynamic Interplay of Anionic Lipids and the Polybasic Region"

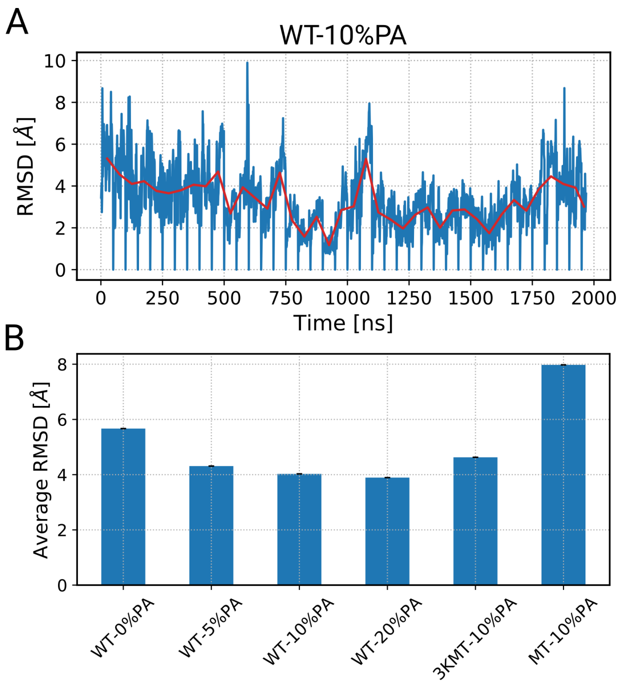


Figure S1: RMSD of the protein. A) RMSD of the protein backbone in slices of 50ns (taking the structure at the beginning of each slice as a reference structure. The running average for each block is shown in red line. B) The average RMSD of the protein backbone for each system. The averaging is done over the RMSD values of all slices considering the data of the second half each slice.

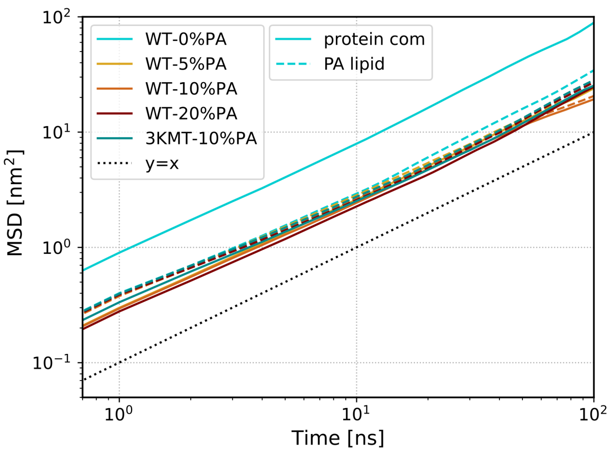


Figure S2: Protein and lipids show similar dynamics. MSD of the COM of the protein as well as P atoms of PAs are represented for different systems.

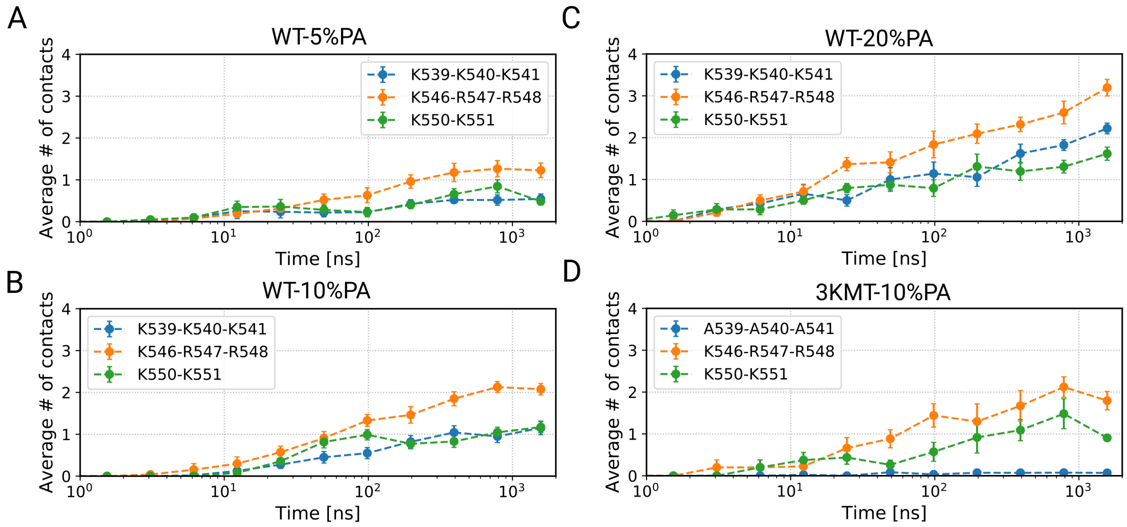


Figure S3: Interaction of PA with different blocks of residues in the polybasic region. The average number of PA lipids bound to the three groups of positively-charged residues throughout the simulation time.

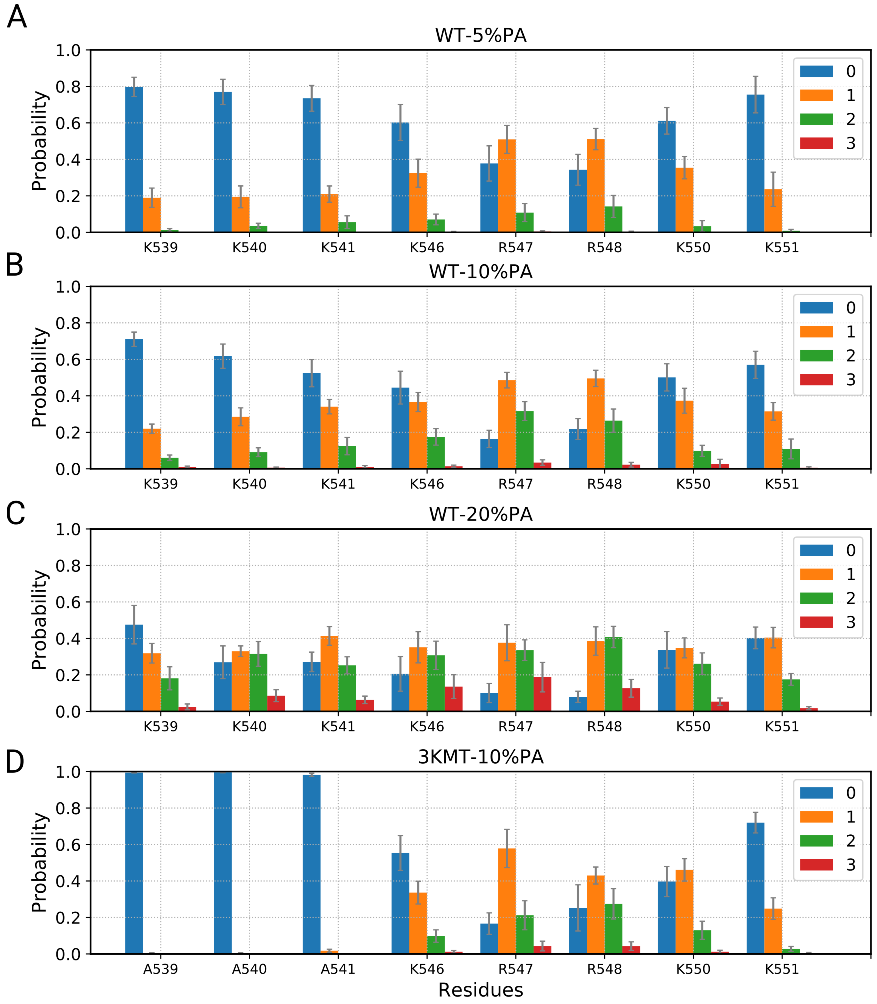


Figure S4: Probability of PA contacts with each proteins' polar residue. The probability of the average number of PA lipid contacts interacting with each residue in the poly-basic region.

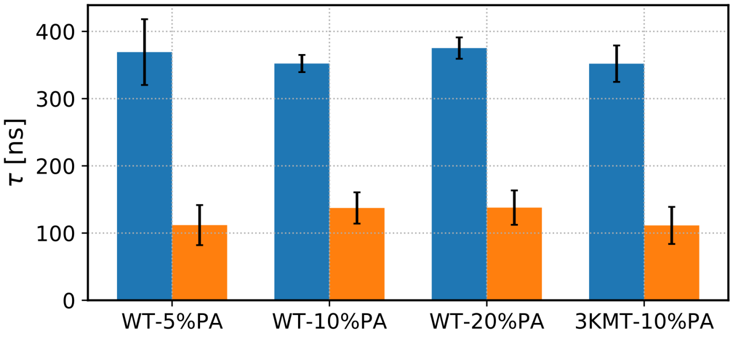


Figure S5: Residence time of PA with the protein. The average residence time of PA molecules with the whole protein (blue) and the polar residues in the poly-basic region (orange). The residence times were calculated from the auto-correlation function of the time series of the number of the number of contacts of PAs with the whole protein and the polar residues.

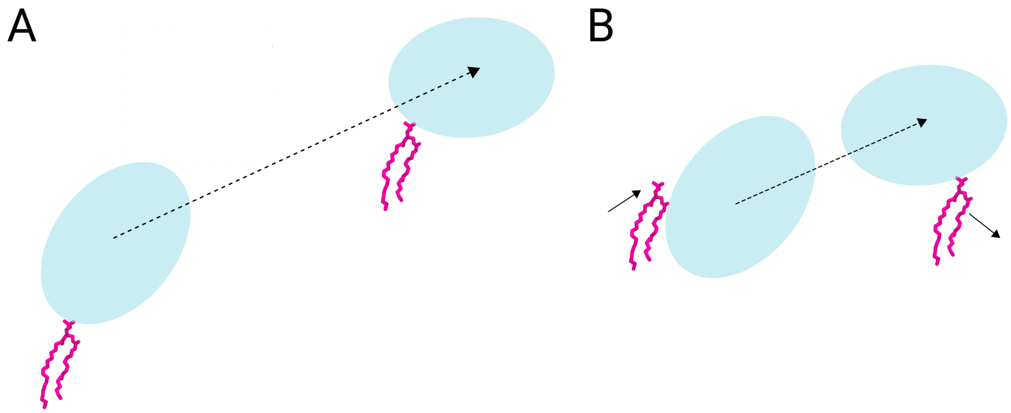


Figure S6: Two scenarios for the protein-lipid binding. Schematic representation of two different scenarios concerning the interaction of a PA molecule with the protein. A) A scenario in which an interacting PA remains bonded to the protein when the protein moves larges distances is shown. B) A scenario in which a PA molecule interacts temporarily with the protein, when the protein traverse a distance in the order of its area is represented. The PA molecule is shown in licorice representation, whereas the protein is depicted via ellipses. The movement of the protein is shown by a dashed line arrow.

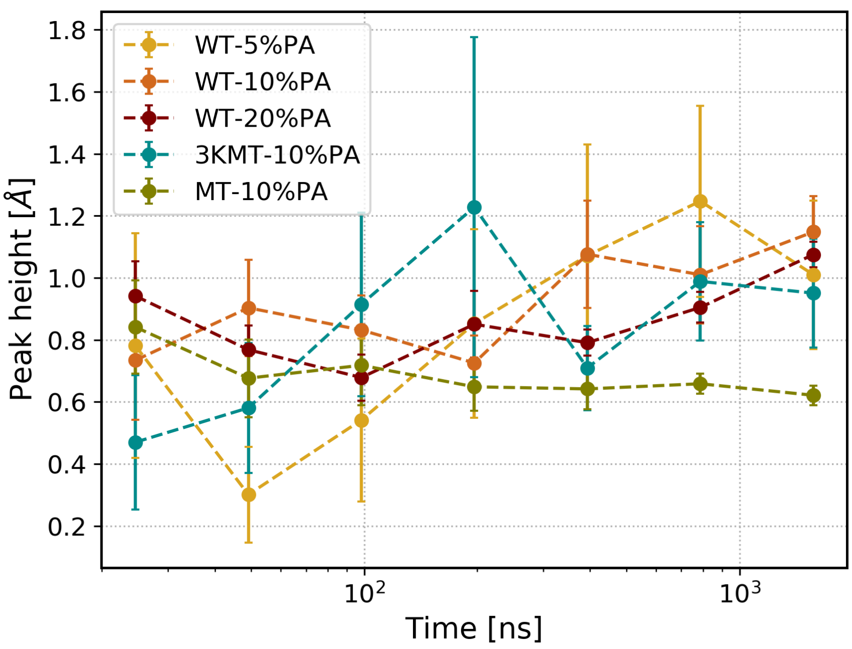


Figure S7: Evolution of the RDF peak during the simulation time. The average of the height of the first peak in the RDF profiles in the course of the simulation.
